# Host genomic variation shapes gut microbiome diversity in threespine stickleback fish

**DOI:** 10.1101/2022.11.14.516485

**Authors:** Clayton M. Small, Emily A. Beck, Mark C. Currey, Hannah F. Tavalire, Susan Bassham, William A. Cresko

## Abstract

Variation among host-associated microbiomes is well documented across species, populations, and individuals. However, the relative importance of host genetic differences for microbiome variation has been difficult to study. While research in humans has shown that genetic variation influences microbiome differences, confounding environmental effects have been very difficult to control. Isogenic laboratory models help isolate host genetic variants to test for influences of the environment or large-effect mutations on the microbiome, but such studies seldom incorporate natural genetic variation. Thus, although many important factors potentially impacting the microbiome have been identified, most studies have failed to test for the direct influence of natural host-genetic variation on microbiome differences within a controlled environment. Understanding the relationship between host-genetic and microbial variation also has biomedical implications, such as understanding why some humans are more susceptible to chronic inflammatory disorders like Crohn’s Disease and Ulcerative Colitis. To directly assess the relationship between host-genetic variation and microbiome variation, we performed a common garden experiment using laboratory lines of genetically divergent populations of threespine stickleback fish – a species that is an outbred model organism commonly used for determining the genetic basis of complex traits in the context of natural genetic variation. Using germ-free derivation and the powerful common garden design with these divergent lines, as well as hybrids between them, we detected clear associations between stickleback genetic dissimilarity and microbiome dissimilarity. Using genome-wide RAD-seq data we also identified regions of the genome underlying differences in microbiome composition. Importantly, we highlight that heritable morphological traits such as body size that are correlated with microbiome dissimilarity also need consideration in future microbiome studies.

## INTRODUCTION

Multicellular organisms harbor a diverse array of microbes on and in their bodies^1–3^. Mutualistic or commensal relationships between hosts and their microbiota are common, with microbes helping to maintain host-health as well as prevent pathogens from successfully colonizing^4,5^. Whether a particular microbe is symbiotic or pathogenic can be context dependent. Chronic inflammatory responses can occur if the host’s immune system recognizes a common symbiont as a pathogen, leading to common diseases including Inflammatory Bowel Diseases (IBD)^6,7^.

Substantial variation in microbiome composition has been documented among host species, populations, and individuals^8–11^. Still, the genetic and molecular mechanisms underpinning these differences have been difficult to study. One particularly difficult area is the influence of host genetic variation on microbiome variation; a relationship essential to understanding population specific disease states and co-evolutionary dynamics between host populations and those of their resident microbes^12–14^. A specific question, for example, is whether greater genomic dissimilarity among individual hosts of the same species leads to greater dissimilarity in their microbiomes^15,16^. Disentangling confounding factors from host genetic influence on the microbiome has proven challenging. In most Genome Wide Associations (GWA) or twin studies, covariance between host genetic and environmental factors cannot be avoided^16–20^.

Although it is easier to control environmental factors in conventional laboratory models, there is often limited genetic variation, which cannot capture what is observed in natural populations^21,22^. Furthermore, human GWA studies have also been biased by limited sampling of global populations that fails to completely sample host genetic diversity^23,24^. What is needed are studies that leverage the standing genetic variation of natural populations of plants and animals that are amenable to controlled laboratory studies. Such research can exploit existing genomic tools while mitigating the limitations described above.

Threespine stickleback fish (*Gasterosteus aculeatus*) is an exceptional model for studying natural genetic variation contributing to complex phenotypes, and has recently been developed as a model for host-microbe interactions through the development of tools, such as gnotobiosis, for microbial experiments^25–31^. Previous studies have focused either on correlating variation in the microbiome of wild stickleback populations with differences in natural environments, or have directly manipulated microbiota in the laboratory^25–28,30^. Despite this progress in stickleback host-microbiome research, a gap remains in that these previous studies were unable to directly assess the relative contributions of the environment and host-genetic variation to microbiome variation.

Here we fill this gap by using genetically divergent, laboratory-raised populations of threespine stickleback fish - and hybrids between these populations - in a controlled and replicated common garden experiment (Figure1; Figure S1). We find evidence of a causal relationship between host genetic and microbiome variation. We also find that the strength of association between host genetic variation and microbiome attributes varies regionally across the genome, providing insight into the genomic architecture of the gut microbiome as a complex trait. Importantly we characterize this relationship in a broader phenotypic context, and document that at least some of the genetic variation associated with microbiome variation is also related to body size. Because such intermediate morphological traits are subject to host genetic variation, they should be accounted for and interpreted accordingly in future studies of host-associated microbiomes.

## MATERIALS AND METHODS

### Fish Husbandry

We generated eight families of threespine stickleback (*Gasterosteus aculeatus*) fish derived from wild-caught Alaskan populations previously maintained in the laboratory for at least ten generations, but with periodic within-population outcrossing. These included three families originating from the freshwater population Boot Lake (N 61.7167, W 149.1167), three families from the anadromous population Rabbit Slough (N 61.5595, W149.2583), and two F1 hybrid families generated from Rabbit Slough females and Boot Lake males (Figures 1 and S1). All experimental families were generated on the same day in a 2-hour period via *in vitro* fertilization. Embryos were incubated overnight in 0.1 µm filter-sterilized antibiotic medium containing 1µl/mL Ampicillin, 0.1µl/mL Kanamycin, 0.312 µl/mL Amphotericin, and 4 parts per thousand (ppt)artificial seawater (Instant Ocean) (Spectrum Brands, Blacksburg, VA, USA). Fin clips from the parents were saved and flash frozen in liquid nitrogen (LN2) to be used later for DNA based parentage assignments.

**Figure 1.**
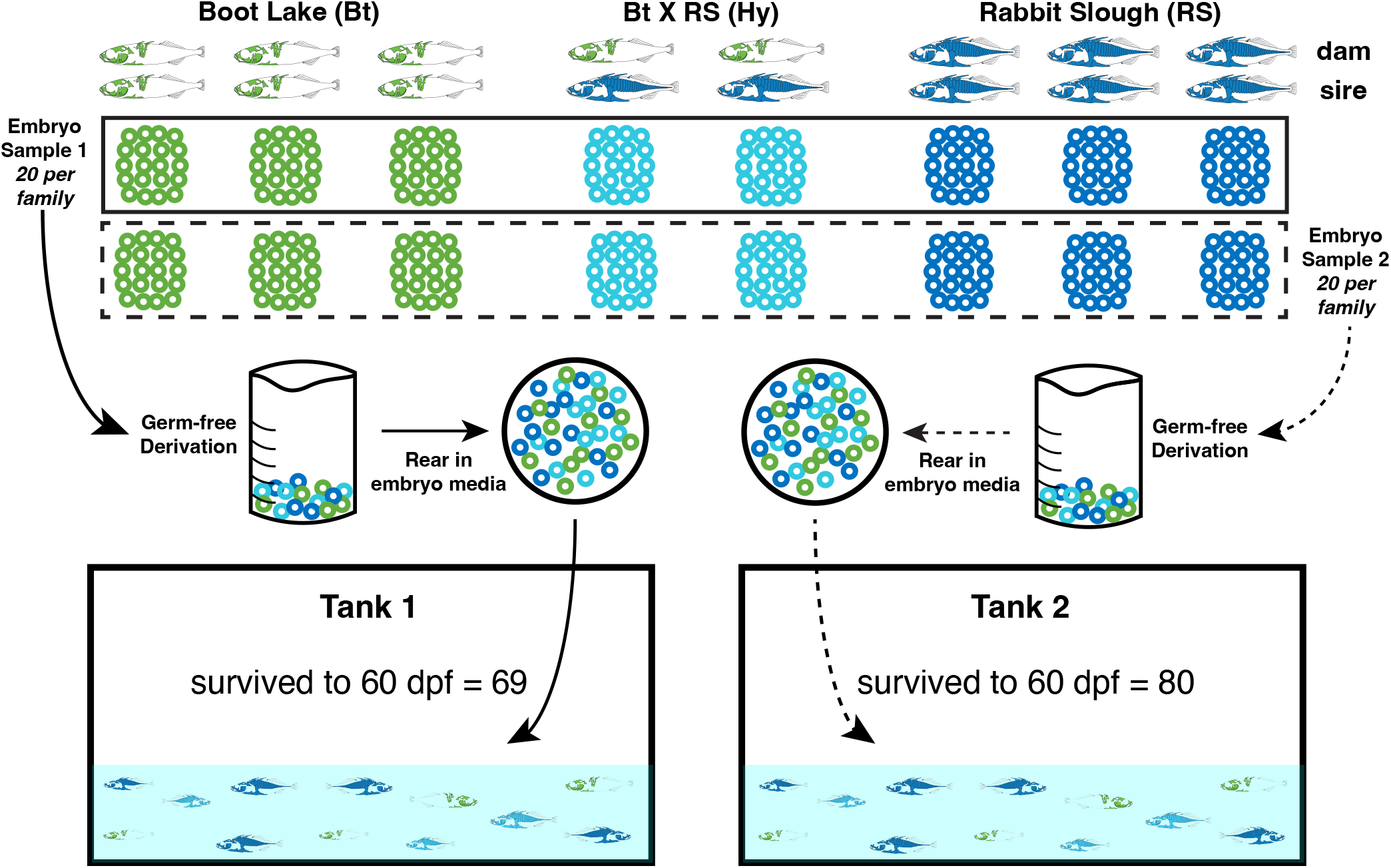
A replicated common garden experimental design enables the controlled measurement of host genetic and environmental influences on the stickleback gut microbiome. We performed eight total stickleback crosses from freshwater (Boot Lake) and oceanic (Rabbit Slough) laboratory populations, including two between-population (“Hybrid”) crosses. We randomly assigned 40, initially germ-free progeny from each cross to two replicate tanks (20 progeny per family, per tank), and we raised the fish to 60 days post fertilization after a 7-day rearing period in two large petri dishes containing non-sterile embryo media. For the 149 surviving fish we determined family membership by parentage analysis using RAD-seq genotypes, and we profiled their gut microbiomes using 16S rRNA amplicon sequencing. The two tank-assigned samples are represented here by solid and dashed lines, and the three cross types (Bt, RS, and Hy) are represented by a respective green, blue, and turquoise color scheme, all of which remain consistent in figures throughout this article.

### Common Garden Experimental Design

Embryos (n = 40) from each family (n = 8) were randomly assigned to one of two groups (n = 20) and combined to comprise two replicate pools of 160 embryos (Figure 1). To remove microbes from the fertilization process and prevent vertical transfer, the two embryo pools were surface sterilized using vacuum-filtered (0.1 µm) 0.2% Polyvinylpyrolidone-iodine (PVPI) and 0.003% bleach diluted in sterile embryo media^25^, maintained in large petri dishes in an incubator (20° C) for ten days, then transferred to two 20-gallon tanks on a recirculating water system. At 60 days post fertilization (dpf) all fish were removed from the tanks for dissection following procedures to standardize the dissecting process (Supplementary Methods). An empty Eppendorf tube accompanied the samples throughout processing and was later used as a negative control to capture environmental microbial contamination. Some mortality is common in clutches of stickleback embryos, and at the end of rearing, tank replicate One contained 69 surviving fish while tank replicate Two contained 80 surviving fish.

### Dissections and Sample Preparation

After euthanasia the offspring were imaged using confocal microscopy to capture phenotypic variation including standard length, and tail clipped for DNA-based parentage assignment. Tail clips were flash frozen in LN2 and stored at -80° C. Guts were dissected following sterile procedures and tracked to account for introduced microbes (Supplementary Methods). Guts were then flash frozen in 1.5 mL Eppendorf tubes previously sterilized via UV irradiation in a Stratlinker (Stratagene, San Diego, CA, USA) at 1000 Joules/cm^2^, and stored at -80° C.

### DNA isolation

DNA was extracted from tail clips from offspring and parents using the Qiagen DNeasy Blood and Tissue Kit (Qiagen, Valencia, CA, USA), and quantified using the Qubit fluorometer broad range kit (Thermofisher, Waltham, MA, USA). Sex was determined using PCR amplification of the male specific GA1 region^32^. Gut samples were homogenized for DNA isolation following a two-step homogenization pipeline (Supplementary Methods), and DNA was then extracted following the Qiagen DNeasy spin column protocol described by Small et al. 2019^29^.

### Library preparation

Genomic DNA from each tail clip was standardized to 10 ng/µL and digested with the restriction endonuclease SbfI-HF (New England Biolabs, Ipswitch, MA, USA), and used to generate RAD-seq libraries^33–35^. The uniquely barcoded samples were multiplexed and run in one lane of sequencing on the Illumina HiSeq 4000 to obtain single-end 150 nucleotide (nt) reads. 12 samples failed during the first round of sequencing and were re-sequenced later in a separate Illumina run following identical methods.

Gut DNA isolations were standardized to 20ng/µl and submitted to the University of Oregon Genomics Core Facility (GC3F) for 16S rRNA (V4 region) amplicon library construction and sequencing (paired-end 150 nt reads) in a single lane on an Illumina HiSeq 4000 (See Supplementary Methods and Small et al. 2019)^29^. Also sequenced were three negative control libraries, generated from a no-tissue extraction sample and two no-PCR-template reactions.

### Genotyping and Parentage Assignment

Tail clip sequencing produced approximately 1.5 million reads per fish from the HiSeq 4000. Raw sequence data were demultiplexed by barcode and filtered using the *process_radtags* program in the Stacks suite^36,37^. Retained reads were then aligned using GSNAP^38^ to the stickleback reference genome (version BROAD S1) obtained from Ensembl. Genotypes were called using the *ref_map* pipeline of the Stacks suite^36,37^. Filtering was then performed using the *Populations* package in Stack*s* with a minimum minor allele frequency of 0.05, a minimum stack depth of 10, and requiring that data be available in 92% of individuals. The resulting 2400 SNPs were then used for parentage assignment by the maximum likelihood program COLONY version 2.0.6.2^39^. COLONY was run using default settings with allele frequencies set to calculate from data and an error rate of 0.001.

### ASV calling, filtering, and enumeration

We demultiplexed 16S reads and performed quality filtering, merging, denoising, and taxonomy classification using tools from QIIME 2 v2018.8.0^40^, generally according to the methods described in Small et al. 2019^29^. Briefly, we used *demux* (*emp-paired*), *vsearch* (*join-pairs*), *quality-filter* (*q-score-joined*), and *deblur* (*denoise-16S*) to define amplicon sequence variants (ASVs). To assign taxonomy to the ASVs we used *feature-classifier* (*extract reads* and *fit-classifier-naive-bayes*) to train a classifier based on the GreenGenes 13_8 99%-clustered OTU database, followed by application of the classifier using *feature-classifier* (*classify-sklearn*). We filtered out any remaining ASVs of mitochondrial or chloroplast origin using *taxa* (*filter-table*). The ASV count tables and ASV sequences were then exported for further analysis using version 3.6.1 of the R statistical language^41^.

Prior to further analysis we filtered any individual 16S libraries that had particularly low sequencing throughput as well as ASVs with particularly low representation among samples or those suspected to be from contaminating (non-fish) sources. Specifically, we excluded one ASV (a likely reagent contaminant) that was in high abundance in the no-tissue negative control library, and we excluded ASVs that were present in only one fish library. We also excluded from downstream analysis individual libraries with fewer than 40,000 total ASV counts. In the case of subsequent multivariate and random forest analyses, ASV count data were normalized to account for depth differences among libraries using the *cpm* function (log=TRUE, prior.count=0.5) from the edgeR R package^42^, whereas the raw counts were used for differential ASV abundance analysis (see below).

### Multivariate community analyses

We visualized stickleback gut microbiomes in community space (defined by 16S ASV abundances) using Principal Coordinates Analysis (PCoA) based on Bray-Curtis dissimilarity, as implemented via the R package phyloseq^43^. We also performed PCoA based on a phylogenetic metric of community dissimilarity, weighted UniFrac^44^. To obtain the weighted UniFrac dissimilarity matrix we aligned ASV sequences using *AlignSeqs* from the R package DECIPHER^45^, optimized parameters for a phylogenetic tree via maximum likelihood using *pml* and *optim*.*pml* functions from the R package phangorn^46^ and generated the weighted UniFrac matrix with phyloseq’s *distance* function. We performed all visualizations and multivariate statistical tests in parallel, using both Bray-Curtis and weighted UniFrac dissimilarity matrices.

To test hypotheses about contributions of experimental design factors to differences in gut microbiome composition among fish we first evaluated a series of permutational multivariate analysis of variance (PERMANOVA) models^47^ using the *adonis2* function from the R package vegan^48^. Both visualization and full, strictly additive models including host population, tank, standard length, sex, and dissector identity as explanatory variables suggested negligible influence from sex and dissector, so we tested fully factorial models excluding these two terms for our primary inferences. Due to the family structure (within populations) inherent in our experimental design, we also performed nested PERMANOVA using the *nested*.*npmanova* function from the BiodiversityR package^49^ to explicitly test population-level effects on microbiome dissimilarity. Last, we also performed tests to evaluate whether the different host genotypic classes (at both population and family levels) in the experiment showed different degrees of community dispersion (i.e. beta diversity) using vegan’s *betadisper* function.

### Random forest classification of host genotype from microbiome data

To test whether gut microbiome structure as quantified by ASV relative abundances provides any predictive potential with respect to host population of origin, we built three random forest (RF) classifiers. The first was trained based on ASV data from tank 1 individuals and applied to tank 2 individuals, the second was based on tank 2 training and tank 1 evaluation, and the third was based on a training set of 80 individuals randomly selected regardless of tank, and application to the remaining 54. We implemented RF models using the randomForest R package and function^50^, with the *strata* and *sampsize* arguments set to minimize bias arising from class imbalance during training.

To evaluate whether average classification accuracy of the RF models was higher than chance expectations, we performed 999 permutations for each model, across which population labels for the training set were randomly shuffled. The accuracies obtained from the original models with unshuffled training data were compared to the null distributions of accuracies from the permutations to infer statistical significance.

### Differential ASV abundance analysis using zero-inflated negative binomial mixed models

We tested for differential abundance of individual ASVs with respect to host population, host size (standard length - SL), tank, and their interactions by fitting zero-inflated and standard negative binomial mixed models, implemented via the R package NBZIMM^51^. We deemed ASVs suitable for zero-inflated negative binomial (zinb) models if they had zero counts in at most 80% of the individuals, but in at least 5%. For those ASVs with zeroes comprising less than 5% of the counts we used standard (nb) models. To fit the models, we also included host family as a random effect, and we summarized the fits from the NBZIMM functions *glmm*.*zinb* or *glmm*.*nb*, including tests for model terms, using type-II analysis of variance as implemented by the R package car^52^. P-values across ASVs for each hypothesis test were adjusted to control the False Discovery Rate^53^.

### Genome-wide association of host genetic and microbiome dissimilarity

To understand the relationship between host genetic and gut microbiome dissimilarity in a genome-wide context we conducted standard and partial Mantel tests^54^ with 1000 permutations and based on individual genetic dissimilarity 1. Calculated from genotypes at all 2408 RAD-seq loci, and 2. Calculated from maximally overlapping sliding windows of five RAD-seq loci, along each of the 21 stickle back chromosomes. We conducted standard and partial Mantel tests using the *mantel* and *mantel*.*partial* functions from the vegan R package^48^.

To quantify individual genetic dissimilarity among fish we calculated the fractional quantity of allelic differences between individuals using the *diss*.*dist* function (with percent=TRUE) from the R package poppr^55^. We used community dissimilarity matrices for these tests based on Bray-Curtis and weighted UniFrac metrics, in parallel analyses. To account for the contribution of host size (SL) differences to the relationship between host genetic and gut microbiome dissimilarity we used partial Mantel tests. Size dissimilarity was simply calculated as the absolute value of the difference in standard length (SL) between two individuals, effectively “Manhattan distance.”

## RESULTS

### RAD-seq data enable parentage assignment in a common garden setting

We obtained 16S amplicon sequencing profiles from the guts of 149 juvenile threespine stickleback, including 69 from the first tank replicate and 80 from the second tank replicate (Figure 1). We excluded 15 individuals from further analysis - six from the first and nine from the second tank - due to low sequencing coverage (< 40,000 ASV-assigned reads after removing contamination). Among the remaining 134 individuals, 63 were female and 71 were male, with mean standard length (SL) estimates (with standard error) for the three different “population-level” genetic backgrounds of 17.368 mm (0.253) for Boot Lake, 18.524 mm (0.301) for Rabbit Slough, and 16.753 mm (0.392) for F_1_ hybrids (Figure S1; Sheet S1). We obtained a total of 2,306 ASVs for final analyses, and the mean number of ASV-assigned reads among the 134 individuals was 238,706.4 (SEM = 11,274.01) (Sheet S1).

Using RAD-seq data generated for the 149 progeny and all 16 of the potential parents, we assigned parentage with 100% confidence to all progeny (Sheet S1). One Boot Lake family did not contain any surviving progeny in either tank, so we inferred that this family sustained complete mortality at an early stage of development. This was consistent with our monitoring of extra embryos (siblings of the 40 embryos per clutch included in the experiment) which also sustained 100% mortality early in development. Remaining for analysis were approximately uniformly distributed family sizes for two Boot Lake families, three Rabbit Slough families, and two F_1_ Boot Lake – Rabbit Slough hybrid families (Figure S1).

### Host genotype, host size, and housing environment influence the stickleback gut microbiome

We found that three factors in our experiment reliably influenced overall microbiome dissimilarity among individual fish: population of origin (“genetic background”), host size, and rearing tank (“environment”) (Figure 2). With respect to Bray-Curtis dissimilarity calculated from 2306 ASV relative abundances, population explained the most variation (PERMANOVA: *R*^*2*^ = 0.056; *F*_2,129_ = 4.235; *p* ≤ 0.001), followed by tank (PERMANOVA: *R*^*2*^ = 0.046; *F*_1,129_ = 8.219; p≤ 0.001), and then by standard length (PERMANOVA: *R*^*2*^ = 0.037; *F*_1,129_= 5.564; *p* ≤ 0.001).

**Figure 2.**
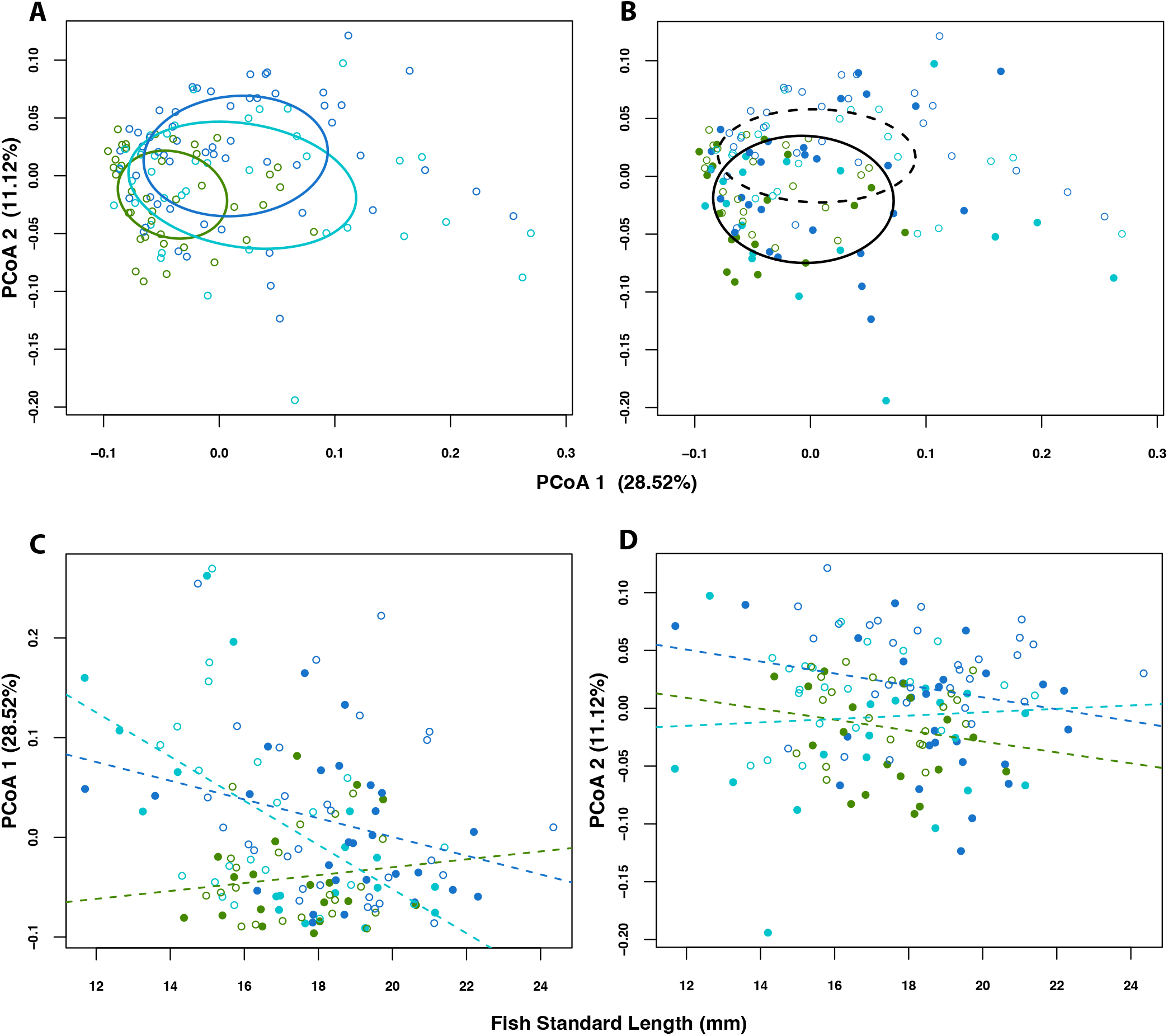
Host genotype (population of origin), host size (standard length), and rearing environment (tank) influence the structure of the stickleback gut microbiome. An ordination of individual stickleback guts in microbial community space using Principal Coordinates Analysis (PCoA), and based on Bray-Curtis Dissimilarity, shows evidence for separation by host family (A) and by tank (B). A scatterplot of values for the first PCoA axis (which explained 28.52% of the total variation in composition) versus fish standard length (SL) shows differing relationships between community structure and fish length among the three different host populations (C). A similar scatterplot, but including PCoA 2 (11.12% of variation explained), suggests a weak relationship between community structure and fish size. In all plots colors represent fish populations (Bt=green, RS=blue, Hy=turquoise). In B-D point style represents rearing tank (tank 1 = closed, tank 2 = open), and in B ellipse line style represents rearing tank (tank 1 = solid, tank 2 = dashed). Ellipses in A-B reflect 95% confidence regions about respective group centroids, and dashed lines in C-D show population-specific slopes from general linear models.

It should also be noted that in a model including first-order interactions (Figure 2) we detected a significant effect of the interaction between population and standard length (PERMANOVA: *R* = 0.028; *F*_2,124_ = 2.148; *p* = 0.007), suggesting that fish size influences the gut microbiome (or vice versa), but to different degrees depending on the host’s genetic background (Figure S2). Beta diversity of the gut microbiome (i.e., dispersion) may differ subtly among the three genetic backgrounds (permutation test: *F*_2,131_ = 3.438; *p* = 0.034), with Boot Lake individuals being less dispersed, on average, than Rabbit Slough or hybrid individuals (Figure 2). This contrasts with other factors including sex, which did not significantly influence overall microbiome dissimilarity, and including the identity of the researcher performing the dissections (Supplementary Methods), which showed a minor effect on overall microbiome dissimilarity (Figure S3).

We found similar trends when analyzing phylogenetic community dissimilarity using weighted UniFrac (Figures S2 and S3). However, standard length explained the most dissimilarity (PERMANOVA: *R*^*2*^ = 0.070; *F*_1,129_= 10.098; *p <=* 0.001), followed by population of origin (PERMANOVA: *R*^*2*^ = 0.034; *F*_1,129_= 2.451; *p* = 0.018), followed by rearing tank (PERMANOVA: *R*^*2*^ = 0.018; *F*_1,129_= 2.624; *p* = 0.046). Again, we noted an effect of interaction between population and standard length (PERMANOVA: *R*^*2*^ = 0.033; *F*_2,124_ = 2.458; *p* = 0.020). We did not find a significant difference in beta diversity among the three genetic backgrounds (permutation test: *F*_2,131_ = 1.345; *p* = 0.282).

To evaluate whether the effect of host genotype in the PERMANOVA models was driven by overall population differences, as opposed to idiosyncratic family effects within populations, we also performed nested analysis based on the Bray-Curtis dissimilarity matrix. We found that population-level variation significantly explained microbiome variation (nested PERMANOVA: *F*_2,127_ = 2.934; *p* = 0.033), relative to family effects (nested PERMANOVA: *F*_4,127_ = 1.194; *p* = 0.160).

### Genomically dissimilar stickleback hosts exhibit more dissimilar gut microbiota

We calculated pairwise genomic dissimilarity among individual fish, based on genotypes from 2,408 RAD-seq loci. Closely related host pairs, on average, tended to have more similar gut microbiomes as measured by Bray-Curtis community dissimilarity (Figure 3). We tested for a statistical association between genomic distance and microbial community distance, while accounting for standard length differences, and found evidence for a subtle but statistically significant positive relationship (Partial Mantel Test: Mantel *r* = 0.0432; *p* = 0.011). We found no evidence for such a relationship between host genomic distance and community dissimilarity as measured by weighted UniFrac (Partial Mantel Test: Mantel *r* = -0.004; *p* = 0.580).

**Figure 3.**
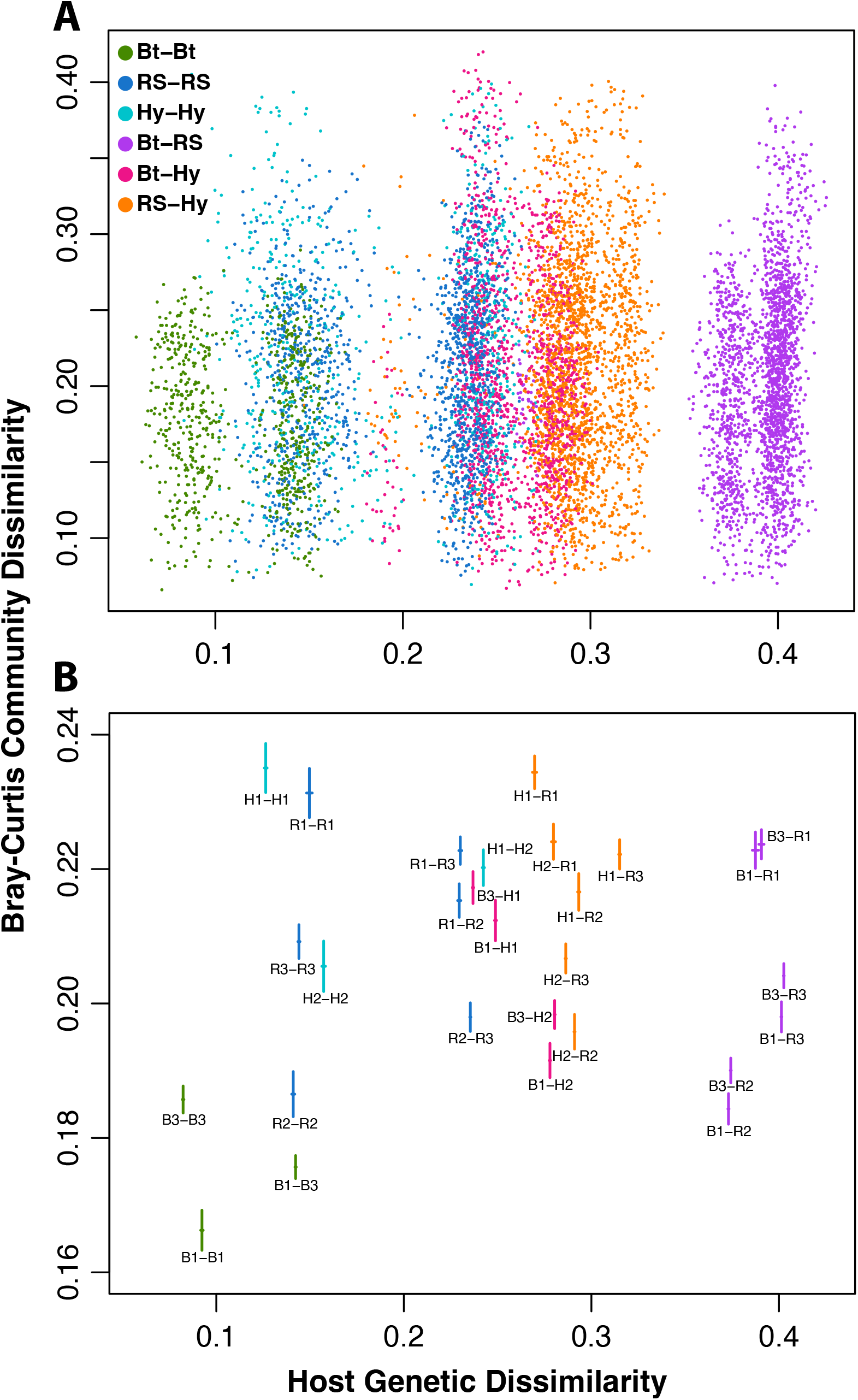
Stickleback that are similar genetically have similar gut microbiomes. A scatterplot (A) of all pairwise Bray-Curtis and genetic (based on 2408 markers) dissimilarity values. Each point is colored (see legend) according to the population combination reflected by the fish pair. A companion plot (B) shows means and means +/- standard errors as crossing lines for family-wise stratification of the pairwise dissimilarities, and each family pair is represented by a text label. Note that the x-axis limits are the same for both A and B, but the y-axis range is smaller for B to better illustrate the distributions of the means.

### Random forest classifiers trained using gut microbiome data predict population membership with less accuracy for fish from hybrid crosses

We applied three random forest models using ASV relative abundances, first by training on fish from Tank 1 to classify population-of-origin for fish from Tank 2, second by training on Tank 2 fish to classify Tank 1 fish, and third by randomly selecting (regardless of tank) 80 fish for training and 54 fish for testing. For two of these models, total accuracy was significantly better than a null distribution generated by permutation (permutation test; p = 0.04, p = 0.073, and p = 0.004, respectively; Figure S4). In all cases the class-wise accuracy was consistently low for F1 (Hy) individuals (0.381, 0.167, 0.333) but higher for Bt (0.429, 0.778, 0.471) and RS individuals (0.517, 0.481, 0.682).

Each model included at least 10 ASVs with relatively high feature importance (Sheet S2), which belonged primarily to Phyla Proteobacteria and Actinobacteria, with representation from diverse classes, including Actinobacteria, Alphaproteobacteria, Betaproteobacteria, Gammaproteobacteria, and Bacilli. One ASV (from Genus *Agrobacterium*) was consistently important for all three models, and several others assigned at least Genus-level taxonomy, including *Luteibacter rhizovicinus, Pseudonocardia halophobica, Sphingomonas, spp*., *Bacillus spp*., and *Edaphobacter spp*., showed high saliency for at least two of the classifiers (Sheet S2).

### Individual bacterial lineages are associated with stickleback genetic background, rearing environment, and genetic-by-environment interaction

We fit negative binomial or zero-inflated negative binomial mixed models to test whether relative abundances of individual ASVs (315 in total) differed among stickleback populations of origin. Among-individual variation was high at the ASV (Figure S5), class (Figure 4), and phylum (Figure S5) levels, but we found statistical evidence of association with population of origin for many ASVs (Sheet S3). Among the 20 ASVs with the largest test statistics (for the effect of population), Class Alphaproteobacteria (Phylum Proteobacteria) dominated, with substantial representation also from Class Chlamydiia (Phylum Chlamydiae). Many (48%) of the ASVs subject to a significant population effect also showed evidence for an effect of interaction between population and tank (Sheet S3). For example, the ASV with the 5th ranking population effect test statistic (assigned to Genus *Bradyrhizobium*) was in low relative abundance among Boot Lake families, but this effect was more pronounced in Tank 2 (Figure S6). Another ASV, assigned to Genus *Agrobacterium* and mentioned above as demonstrating consistent random forest feature importance, was also subject to population and population-by-tank interaction effects but with consistent low abundance in Boot Lake individuals (Figure S6). Other ASVs, for instance one assigned to Genus *Perlucidibaca*, showed strong tank differences without population or population-by-tank interaction effects (Figure S6).

**Figure 4.**
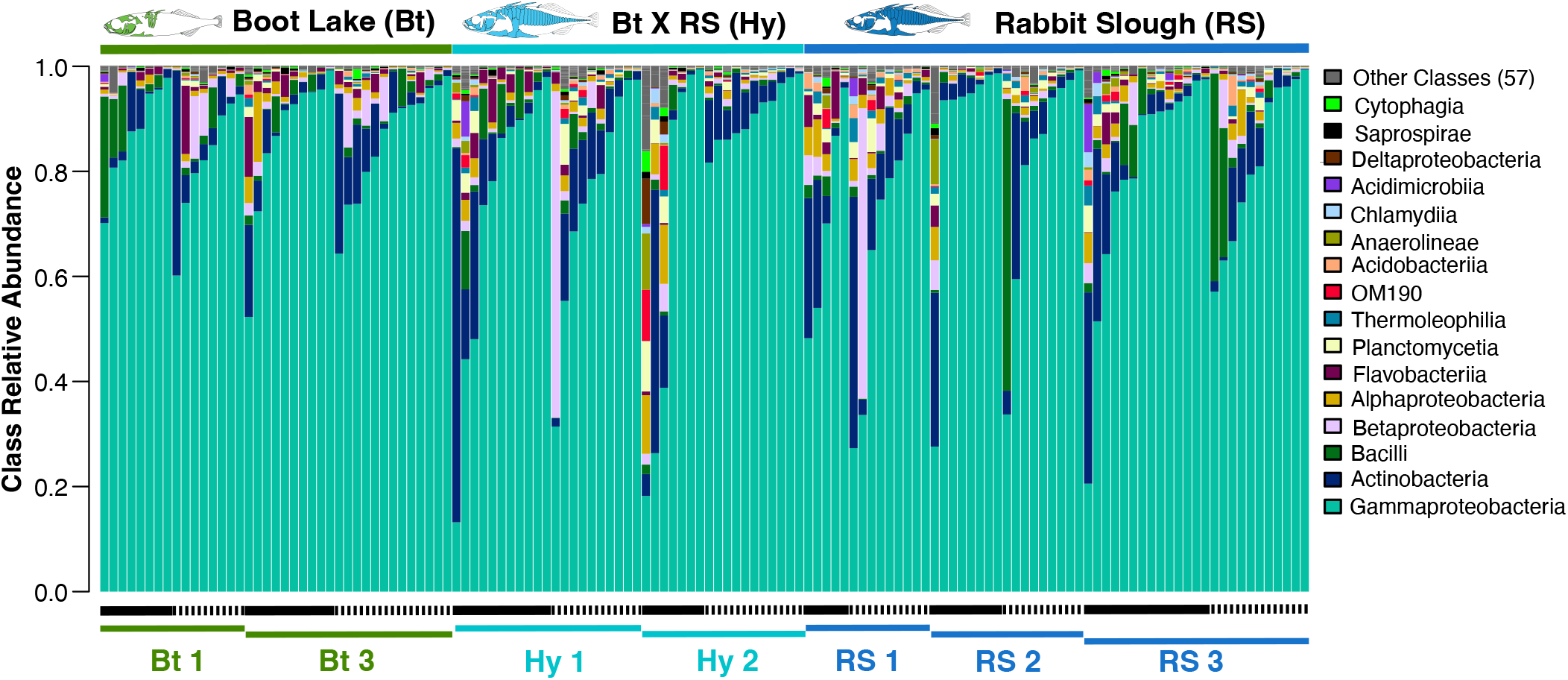
Relative abundance of bacterial classes based on 16S profiling demonstrate substantial variation in community composition among individuals, populations and families, and rearing tanks. Each vertical bar represents the gut microbiome of an individual fish, and bars are ordered within each family-tank combination by increasing abundance of Class Gammaproteobacteria, the most abundant class overall on average. Black horizontal bars below the plot represent tank (Tank 1 = solid, Tank 2 = dashed), and colored horizontal bars represent stickleback families. The legend to the right of the plot provides a key to the 16 most abundant (on average, from bottom to top) classes.

### Gut microbiome and host genetic associations vary in strength and mode along the stickleback genome

Using a sliding window approach using partial Mantel tests, we identified 31 regions of the stickleback genome at which genetic dissimilarity is likely positively associated with gut microbiome (Bray-Curtis) dissimilarity (Figure 5 and Sheet S4). Based on a parallel analysis using Weighted UniFrac to quantify community dissimilarity we identified just six genomic regions (Figure S7), but all except one overlapped with the 31 strongest candidate regions from the Bray-Curtis analysis. These five intersecting candidate regions lie on chromosomes 1, 14, 16, 18, and 20, in marker position intervals of 1.923, 0.406, 3.077, 0.573, and 3.372 Mb, respectively. The strength of the relationship is clear between gut community (Bray-Curtis) and host genetic dissimilarity for these regions (Figures 5 and S8) relative to all markers (Figure 5) and to five randomly sampled regions of the same approximate size (Figure S8).

**Figure 5.**
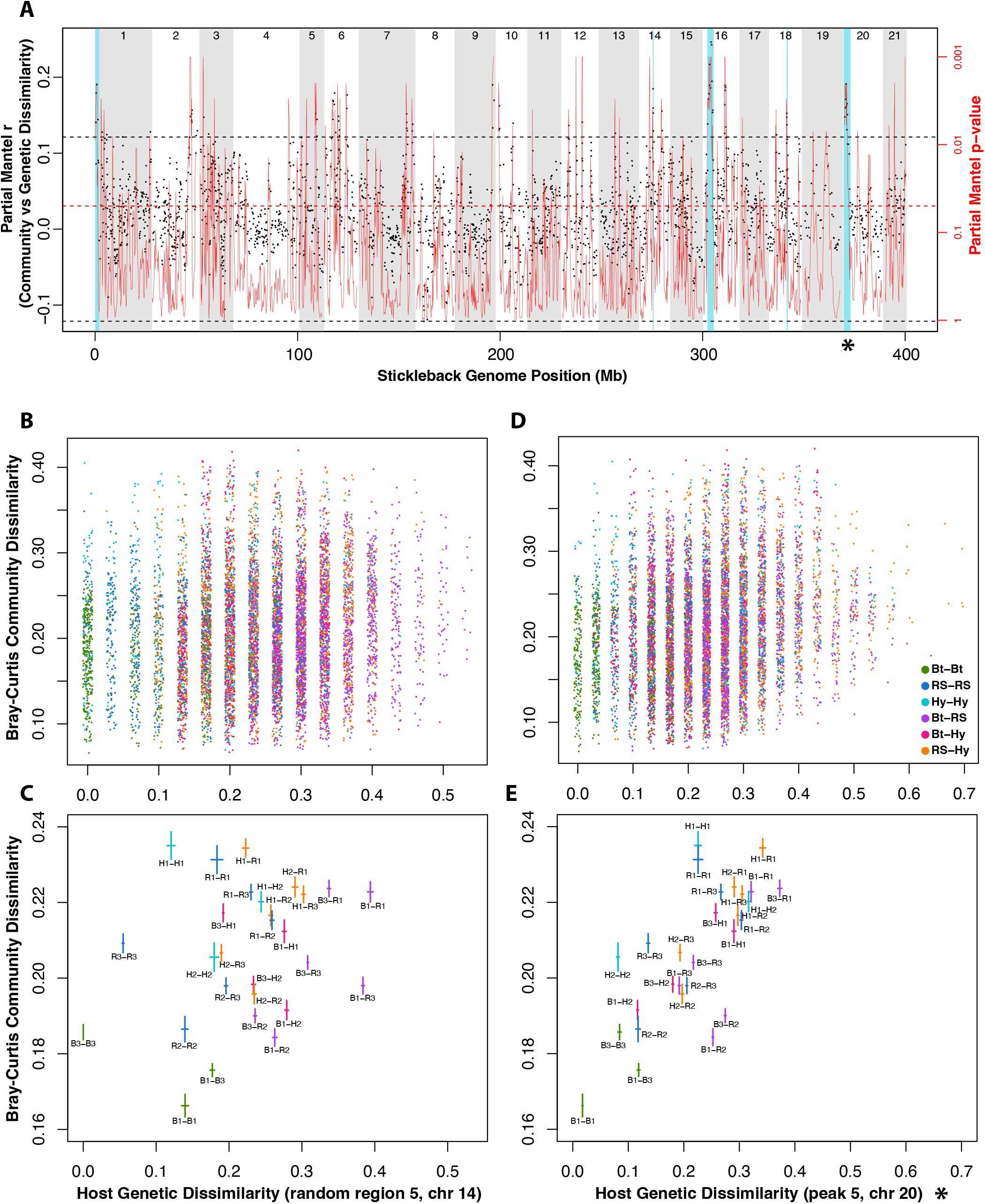
The strength of the relationship between host genetic and gut microbiome dissimilarity varies along the stickleback genome. A sliding window analysis (A) shows the strength of host-genetic vs. gut microbiome dissimilarity associations across chromosomes. For each overlapping window of 5 (RAD-seq) makers, a partial Mantel test was used to assess the relationship between genetic and microbiome (Bray-Curtis) dissimilarity. The left (black) y-axis and points indicate the test statistic, and the right (red) axis and lines represent the lowess-smoothed p-value distribution from the statistical tests. Note that the red p-value y-axes are on a log10 scale, ascending from high to low, to make easier comparisons with the black test statistic y-axes. Points (marker windows) above the top, dashed line represent regions of the genome especially strongly (i.e. beyond null expectations) associated with microbiome differences. The red, dashed line simply indicates a p-value of 0.05. Blue bands show 5 “peaks,” window blocks that are consistently significant across frequency-based (Bray-Curtis) and phylogenetic (Weighted UniFrac) metrics of community dissimilarity. These peaks likely correspond to regions of the stickleback genome that influence gut microbiome differences. Alternating white and gray bands demarcate stickleback chromosomes. Plots of host genetic and gut microbiome dissimilarity for a randomly selected region of 15 markers on chromosome 14 (B-C) show a relatively weak association, whereas the 15 markers in peak 5 (D-E, and marked with an asterisk) show a strong, positive relationship. Each point in B and D are colored (see legend in D) according to the population combination reflected by the fish pair. In Plots B and D points are “jittered” about discrete degrees of genetic dissimilarity (x-axis) to reduce obfuscation. Plots C and E show means and means +/- standard errors as crossing lines representing family-wise stratification of the pairwise dissimilarities, with each family pair represented by a text label.

## DISCUSSION

### Controlled common garden experiments are essential for understanding determinants of animal microbiomes

Several important features of our common garden experimental design are useful in addressing previously challenging questions. We subjected genetically variable individuals to identical rearing conditions, truly co-housed and with replication, allowing the disentanglement of host genotypic from environmental effects on the gut microbiome. This degree of experimental control has been implemented more commonly for plant microbiome studies^56–59^, but it is logistically difficult for studies of free-living animal species. Because animal experiments with unrestricted co-housing require the tracking of individuals via marking or genotyping, a more convenient alternative has been to keep subjects with different genotypes physically separated but generally exposed to similar conditions^28,29,60–63^. In contrast, we incorporated unrestricted co-housing in our current study, an approach taken in only a few other studies^64–66^.

Importantly, external fertilization and early surface sterilization of embryos via gnotobiotic techniques developed for stickleback^25^ allowed us to control for vertical transfer of microbes and to create a common conventional starting point for microbial colonization among individuals. Germ-free derivation has also been leveraged to understand host genetic contributions to microbiome attributes in *Drosophila*^67^, dung beetles^68^, mice^69^, and zebrafish^64^, although some of these studies relied on inbred lines or induced mutations, which places limitations on their ability to recapitulate and interrogate the landscapes of genetic variation often observed in natural populations. The design of our current study included both crucial elements: genetic variation sampled recently from natural stickleback populations coupled with a highly controlled, standardized environment. Additionally, by using genetically distinct populations of threespine stickleback and their F1 progeny, we broadened the continuum of genetic variation along which to assess patterns of inheritance in both regional genomic and quantitative genetic contexts.

### Important interactions among host genetic background, rearing environment, and host morphology in microbiome structure

The primary objective of our study was to test whether host genetic variation determines the composition of the stickleback gut microbiome, and we provide three lines of evidence that support this hypothesis. First, we document clear partitioning by host population of variation in community space. Second, ASV abundance data predicts host population with above-noise accuracy using random forest classifiers. Third, among-individual genetic differences along the host genome accompany microbiome differences. The general strength of this pattern was greater for analyses based on Bray-Curtis dissimilarity relative to those based on weighted UniFrac, suggesting that groups of bacterial lineages with recent shared ancestry may be collectively influenced by host genetics.

Interestingly, we also found that relationships between host genetic variation and gut microbiome variation differed depending on other study variables, namely rearing environment (tank) and fish size (standard length). These statistical interactions between host genotype and other factors are not surprising, as other studies of host-microbiome associations have revealed similar patterns. For example, a tightly controlled, recent study of influences on the stickleback gut microbiome revealed that host genotype effects were more pronounced during infection with the cestode parasite *Schistocephalus solidus*, relative to controls^66^, and authors of a study in which two Japanese quail genotypes received different cholesterol diets found that diet interacts with host genotype to influence the intestinal microbiome^70^.

Our observed connections among stickleback genotype, standard length, and gut microbiome composition highlight an increasing appreciation for the idea that a multitude of heritable host traits likely unrelated to immunobiology *sensu stricto* make significant contributions to microbiota. Indeed, traits such as body size in killifish and cod^71^ and mice^72^, and leaf attributes and physiological traits in *Picea* spruces^73^, have been shown to co-vary with host-associated microbiome attributes, but without necessarily explaining the totality of host genetic contributions to microbiome variation. In the case of our study, we noted a relationship of varying strength between SL and community composition across the three genotypes, and that host genetic variation still explained microbiome variation after accounting for the size differences.

One outstanding question, however, concerns the directionality of potential causal relationships among these variables. Body size or growth function parameters, which show high heritability in stickleback fishes^74,75^, may determine which microbes can colonize and/or persist in the gut. Standard length can vary substantially between adults in threespine stickleback populations and is also often used as a metric for determining developmental stage of juveniles^76^. As our fish were 60 dpf – still juvenile – it is possible differences in developmental rate among genotypes could explain our results.

Alternatively, it is conceivable that a fish’s gut microbiome, determined in part by its genotype, affects metabolism and ultimately growth to dictate size. In any case, consideration of a host’s composite phenotype, including morphological, behavioral, metabolic, and immunological traits, is important when conceptualizing host genetic contributions to the microbiome. Traits like body size within the stickleback and other host systems, for example, should be explicitly evaluated and accounted for whenever possible, when the primary interest of studies (e.g., GWAS) is the identification of host genetic microbiome determinants. Importantly, our ability to document the interactions among these variables further highlights the strength of the common garden experimental design.

### Gut microbiome dissimilarity increases with greater host genetic dissimilarity

One important prediction from our conceptual understanding of host-microbe co-evolutionary dynamics is that continuous genetic differentiation among individual hosts in a population, particularly owing to differing immunogenetic repertoires, will translate to compositional dissimilarity of the microbiome^16,77^. Tests of this prediction for hosts of the same species have been rare, are seldom performed using controlled experiments, and have provided mixed results. For example, an environment-controlled study of *Daphnia galeata* hatched from different sediment layers^61^ tested for an association between genome-wide genetic dissimilarity and microbiome dissimilarity, and it yielded no such relationship. On the other hand, authors of an exome sequencing-based study compared wild-caught house mice from five natural populations and did find evidence for a positive relationship between host genetic and gut microbiome dissimilarity^60^, as did authors of a microsatellite-based study of wild-caught threespine stickleback from six lake populations^26^.

We found a positive relationship between genetic dissimilarity calculated from 2408 RAD-seq loci across the stickleback genome and Bray-Curtis gut microbiome dissimilarity, accounting for differences in fish size. This pattern appears to be driven largely by smaller genomic differences and more similar microbiomes, on average, between Boot Lake individuals, and more generally by smaller genetic and microbiome dissimilarities between individuals within, relative to between, families (Figure 3). Interestingly, community dissimilarity was not greatest for individual pairs with a Boot Lake and Rabbit Sough individual (the most genetically dissimilar pair type in the experiment), suggesting that allelic dominance at host loci may be important for inheritance with respect to microbiome composition.

Not surprisingly, this structural relationship between genetic variation and beta diversity was also borne out by multivariate analyses treating population of origin as a discrete factor. We observed that the gut microbiota of Boot Lake individuals was less dispersed, on average, relative to Rabbit Slough or hybrid individuals. Boot Lake individuals, considered as a group in this experiment, have experienced more inbreeding and exhibit on average lower genetic variation than Rabbit Slough individuals.

Our abilities to predict population membership of fish based on their gut microbiota also varied by genetic diversity within the groups. Random forest classifiers trained using ASV relative abundances were most accurate for fish belonging to pure freshwater (Boot Lake) or pure anadromous (Rabbit Slough) populations. Accuracy declined when predicting membership for F1 individuals from the hybrid crosses. Deeper investigation, including greater sampling of progeny from a more diverse panel of crosses, is needed to better understand what aspects of genetic variation and forms of inheritance (i.e., non-additive) are driving the patterns observed here.

### Host genomic regions contribute differentially to variation in the gut microbiome

Our sliding window analyses of the relationship between host genetic and microbiome dissimilarity suggest that several regions of the stickleback genome contribute large effects to gut microbiome variation among individual fish, relative to the genome average. The distribution of effect sizes for the analysis based on Bray-Curtis community dissimilarity is similar to that for an analysis in which we tested for correspondence between genetic dissimilarity and standard length (SL) differences (Figure S7), suggesting for this metric of community dissimilarity anyway, a comparable genomic architecture in the broad sense.

As was the case for several other analyses, however, measurement of microbiome dissimilarity using weighted UniFrac resulted in fewer large-effect regions of association than analysis based on Bray-Curtis dissimilarity.

Nevertheless, five genomic regions from the two analyses overlap, lending support for at least several substantial genetic effects, each on a different stickleback chromosome. In addition, we controlled for standard length differences via partial Mantel tests, and none of the five regions overlapped with large-effect regions from the SL analysis, so host genetic effects other than those on body size explain the differences in community structure.

The genomic regions of association we identified are large, covering hundreds of kb in some cases, so narrowing intervals to putative causal variants would likely require larger GWA or fine-mapping analyses, with considerably larger sample sizes. At least one other study has included an analysis (QTL mapping) of association between genomic variants and gut microbiome composition in threespine stickleback from two British Columbia lake populations^78^. Although mapping intervals were also wide in that study, there appears to be minimal overlap between QTL for the latent community variables mapped therein and association peaks from our analyses, with the possible exception of overlapping regions on chromosomes 2, 14, and 20.

### Conclusions and perspective

Evidence from our carefully controlled common garden experiment in stickleback inclines us to conclude that host genomic diversity is a definitive facilitator of beta diversity for gut microbiomes among individual fish, and that inheritance of structural microbiome traits is not strictly additive. The replicated tank element of our common garden design also demonstrated that rearing environment is an important factor, and that the interaction between genetic and environmental variation (G-by-E) influences the microbiome as well.

Looking forward several attributes should be prioritized for diverse studies that are devoted to understanding animal host genetic contributions to microbiota. Approaches in which standing genetic variation can be leveraged and measured across the host genome, as compared to gross population or inbred line membership, permit much more nuanced understanding of genomic architecture and possible mechanistic connections between host genes and microbes. In addition, for those studies with inherent reliance on inferences from host genomic data (e.g. GWAS, QTL mapping), efforts should be made to limit the impact of variables that confound genetic effects, through tools like gnotobiotic protocols and unrestricted co-rearing, all of which are possible to implement for powerful outbred fish models such as threespine stickleback.

## Supporting information

Supplementary Sheet 1

Supplementary Sheet 2

Supplementary Sheet 3

Supplementary Sheet 4

## ACKNOWLEDGEMENTS

We thank Maggie Weitzman and the University of Oregon Genomics and Cell Characterization Core Facility (UOGC3F) for advice on, and optimization of, 16S library preparation and sequencing. We also thank Micaela Burns and Natalie Verhoeven for measuring the standard length of each fish. We also thank Jake Searcy for discussions about the data set featured in this article, particularly with respect to machine learning approaches. This work was supported by NSF Grant OPP-2015301 (to C.M.S., S.B., and W.A.C), NIH grants P50GM098911and R24RR032670 (to W.A.C), and a National Institute of Health NRSA fellowship F32GM122419 (to E.A.B). This work was further supported by Oregon Research Excellence funds (to W.A.C) and funds from the UO Office of the Vice President for Research and Innovation through an Incubating Interdisciplinary Initiatives Award (to W.A.C. and E.A.B) and a Faculty Research Award (to E.A.B).

## AUTHOR CONTRIBUTIONS

The study was designed by C.M.S., E.A.B., and W.A.C. Fish husbandry and maintenance were performed by C.M.S. and E.A.B. Dissections and tissue preservation were performed by C.M.S., E.A.B., M.C.C., and S.B. Tissue processing was performed by C.M.S. DNA isolations and library preparations were performed by E.A.B. and M.C.C. Parentage analysis was performed by C.M.S., E.A.B., and H.F.T. Microbiome and statistical analyses were performed by C.M.S. The initial manuscript draft was written by C.M.S. and E.A.B., with subsequent editing and contributions from all authors.

## DATA AVAILABILITY STATEMENT

Raw 16S amplicon sequencing and RAD-seq reads will be available upon peer-reviewed publication in the NCBI SRA. Other data files and code not included with the Supplementary Information will be available upon peer-reviewed publication in Dryad.

## Notes

### Competing Interest Statement

The authors have declared no competing interest.

