## Supplementary Information for "Host genomic variation shapes gut microbiome diversity in threespine stickleback fish"

### SUPPLEMENTARY MATERIALS

#### Supplementary Figures

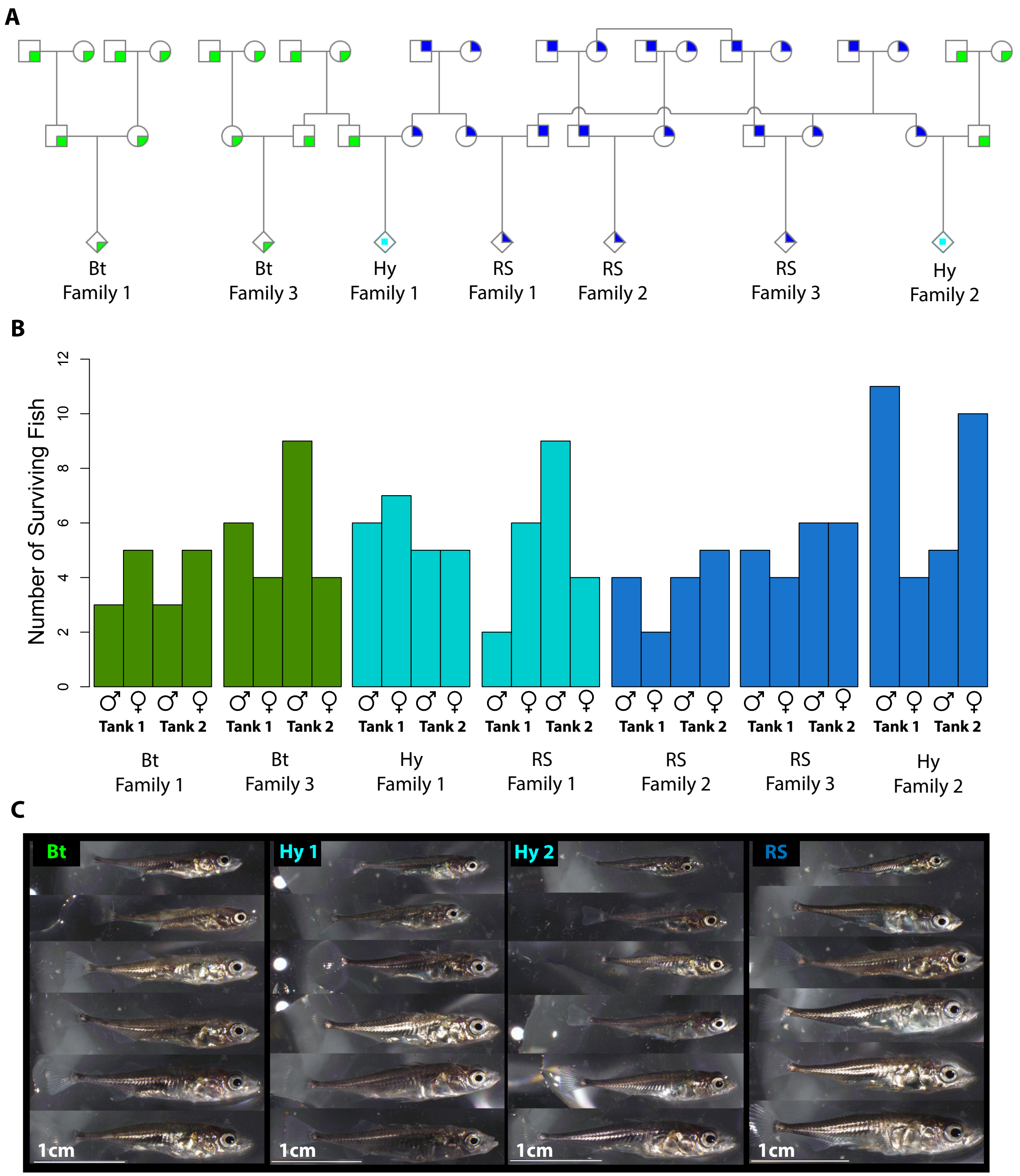


Figure S1. Information from the replicated common garden experiment summarizes representation outcomes. A pedigree (A) shows relatedness among experimental fish going back two generations. Squares and circles are individual ancestors, and diamonds represent the full sibships profiled in this study. Note that Boot Lake (Bt) Family 2 is not included, as none of the embryos from that cross (family) survived (see Methods). The number of surviving fish for each sex, family, and tank (B) show the distribution of individuals for factors of interest in our analysis. A random selection of experimental fish images (C) shows gross variation in size and morphological features among all fish in the study. Represented are 6 Bt (Freshwater) fish, 6 RS (Oceanic) fish, and 12 Hy (F1) fish.

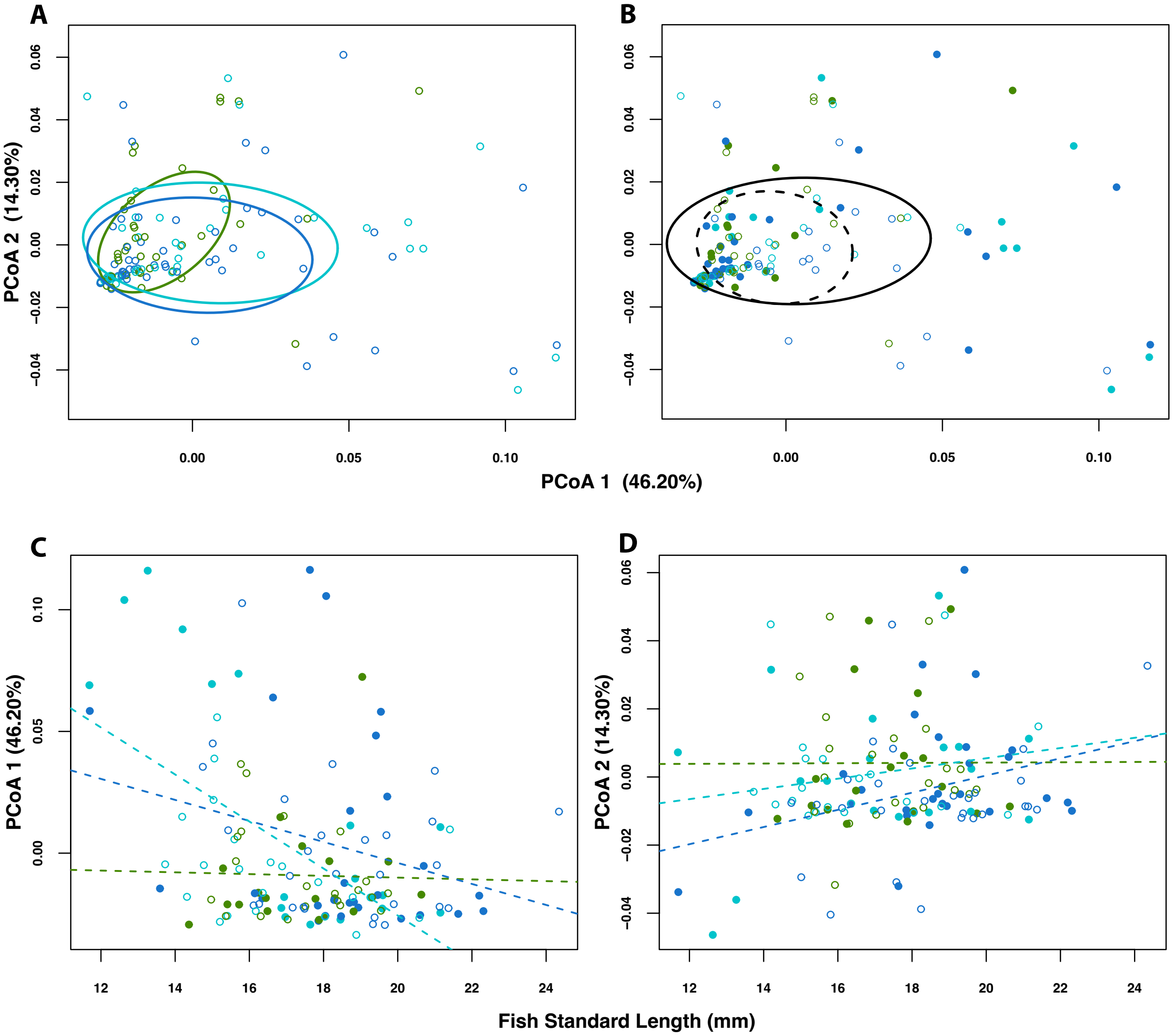


Figure S2. Host genotype (population of origin), host size (standard length), and rearing environment (tank) variably influence the structure of the stickleback gut microbiome. An ordination of individual stickleback guts in microbial community space using Principal Coordinates Analysis (PCoA), and based on Weighted UniFrac, shows subtle evidence for separation by host family (A) but minimal evidence for a tank effect (B). A scatterplot of values for the first PCoA axis (which explained 46.20% of the total variation in composition) versus fish standard length (SL) shows differing relationships between community structure and fish length among the three different host populations (C). A similar scatterplot, but including PCoA 2 (14.30% of variation explained), suggests a weaker relationship between community structure and fish size. In all plots colors represent fish populations (Bt=green, RS=blue, Hy=turquoise). In B-D point style represents rearing tank (tank 1 = closed, tank 2 = open), and in B ellipse line style represents rearing tank (tank 1 = solid, tank 2 = dashed). Ellipses in A-B reflect 95% confidence regions about respective group centroids, and dashed lines in C-D show population-specific slopes from general linear models.

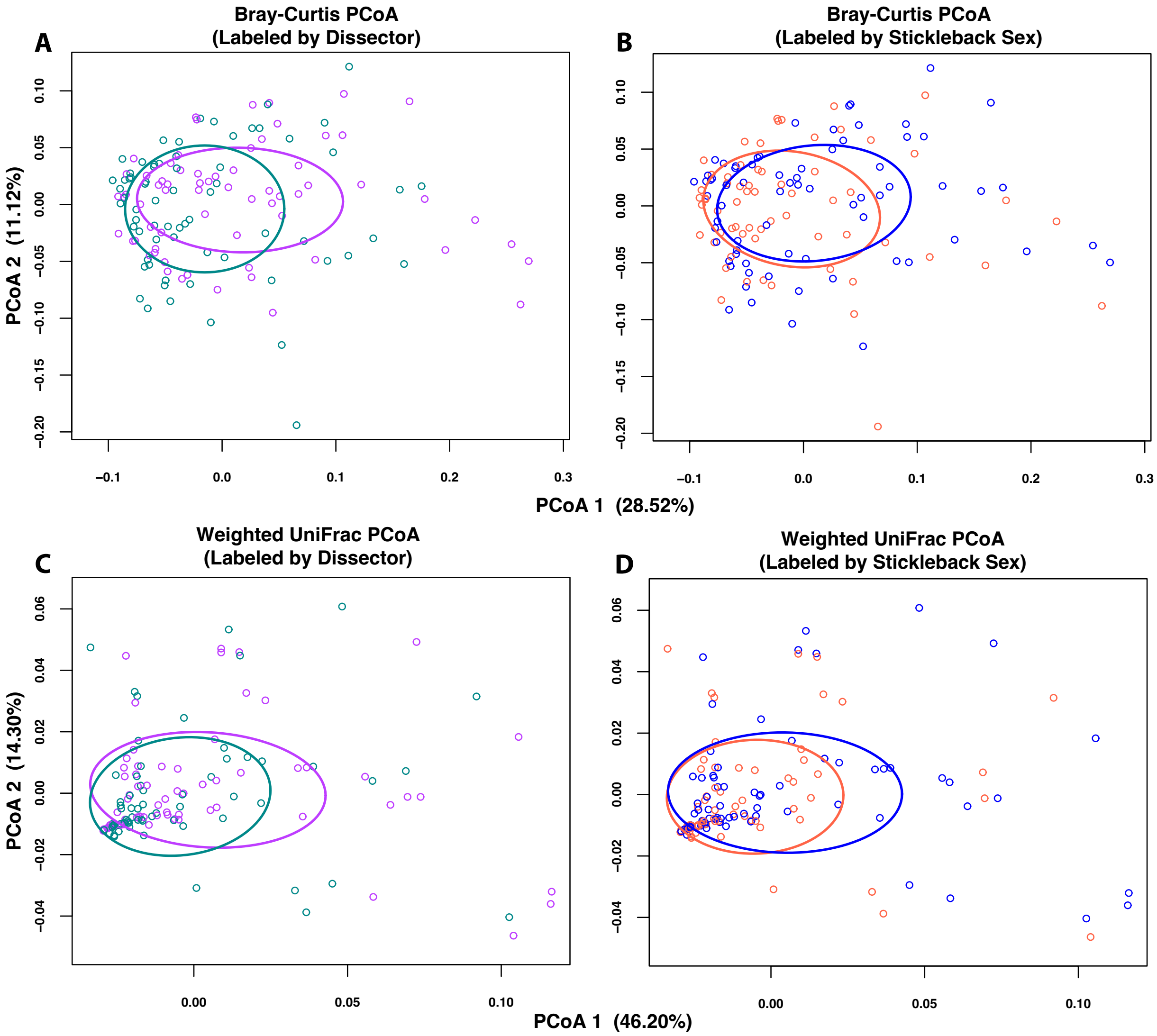


Figure S3. Fish dissector and fish sex do not contribute appreciably to variation in the measured stickleback gut microbiome. Ordinations of individual stickleback guts in microbial community space using Principal Coordinates Analysis (PCoA) based on Bray-Curtis Dissimilarity (A-B), or based on Weighted UniFrac (C-D), show little to no evidence for separation by the person performing the gut dissection (A and C), or by the sex of the fish (B and D). In A and C colors represent dissector (dissector 1 = pink, dissector 2 = teal), and in B and D colors represent stickleback genotyped sex (female = red, male = blue). Ellipses reflect 95% confidence regions about respective group centroids.

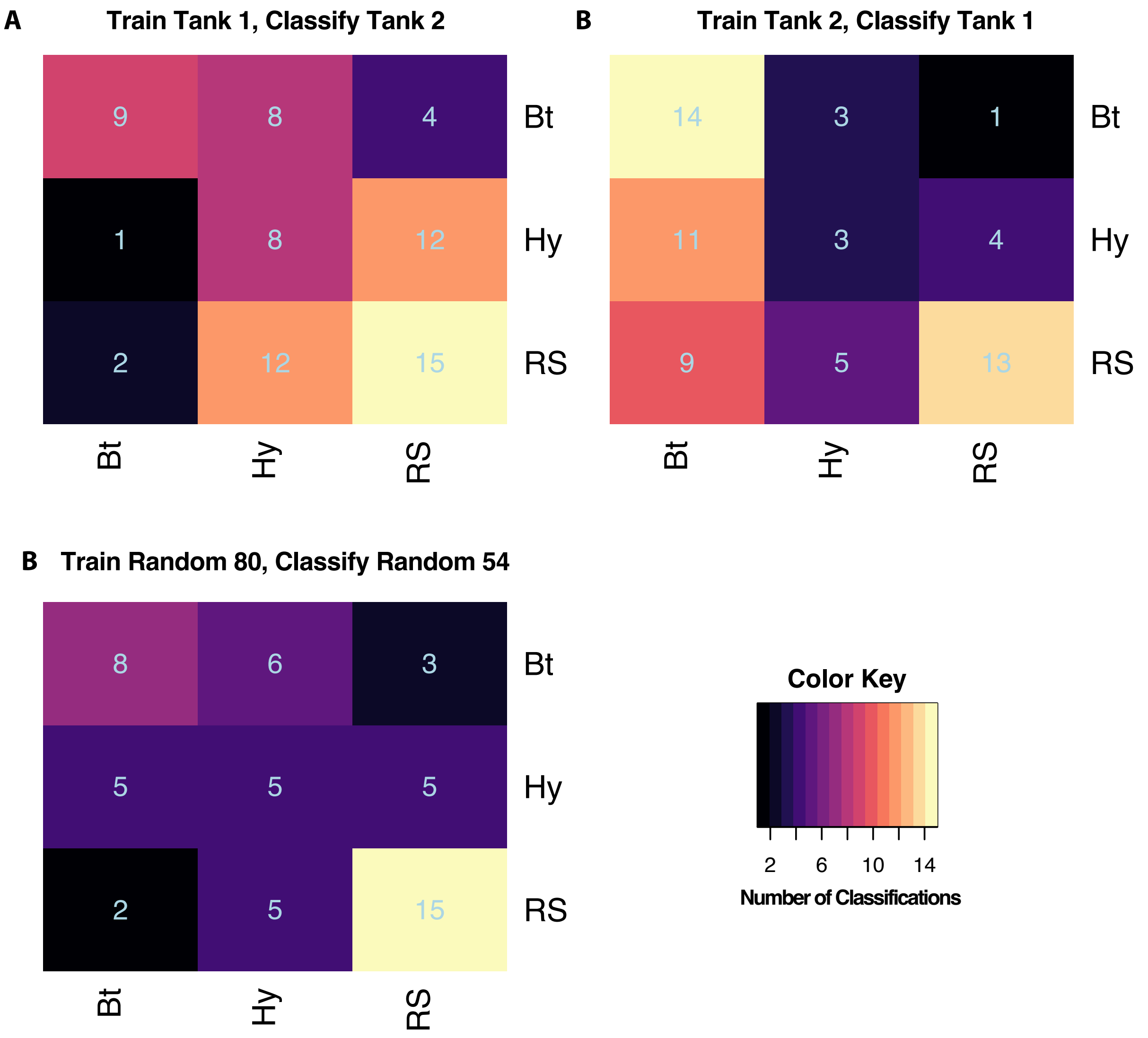


Figure S4. Random forest classifiers predict, with minimal but better-than-random accuracy, stickleback population of origin from gut 16S data alone. Confusion matrices for random forest models trained using Tank 1 fish and tested on Tank 2 fish (A), trained using Tank 2 fish and tested on Tank 1 fish (B), and trained using 80 randomly selected (regardless of tank) fish and tested using the remaining 54 fish (C). Classifications are listed row-wise in each matrix. For example, in A, 9 Bt fish were accurately classified as Bt, 8 Bt fish were erroneously classified as Hy, and 4 Bt fish were erroneously classified as RS. Note that accuracy for classification of Hy fish is consistently lower than accuracy for Bt and RS fish.

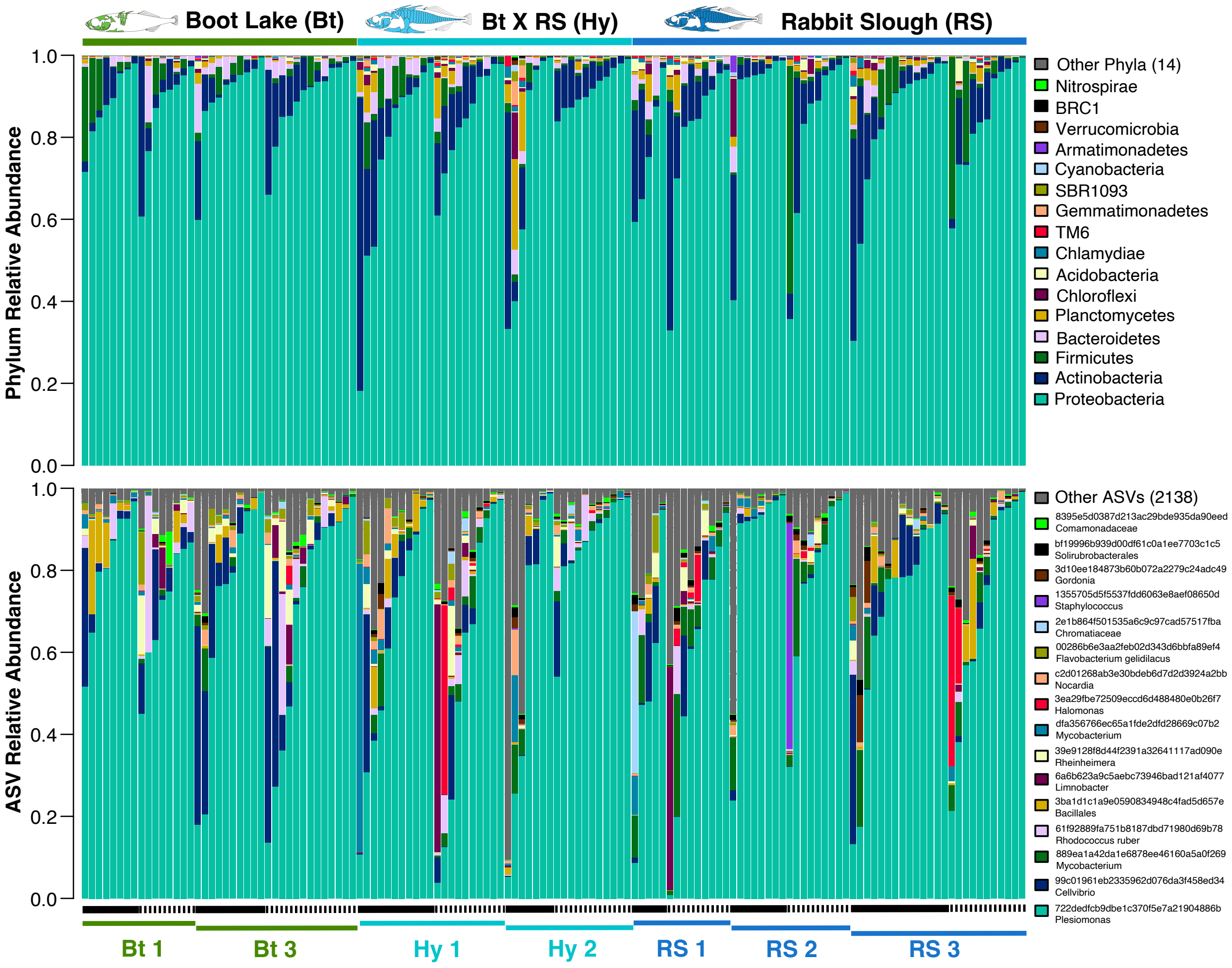


Figure S5. Relative abundance of bacterial phyla (A) and individual ASVs (B) based on 16S profiling demonstrate substantial variation in community composition among individuals, populations and families, and rearing tanks. Each vertical bar represents the gut microbiome of an individual fish, and bars are ordered within each family-tank combination by increasing abundance of Phylum Proteobacteria (A), and (Plesiomonas spp.), ASV 722dedfcb9dbe1c370f5e7a21904886b (B), the most abundant taxa, respectively. Black horizontal bars below the plots represent tank (Tank 1 = solid, Tank 2 = dashed), and colored horizontal bars represent stickleback families. The legends to the right of the plots provide a keys to the 16 most abundant (on average, from bottom to top) phyla and ASVs. For ASVs, the highest level of classified taxonomic resolution is listed.

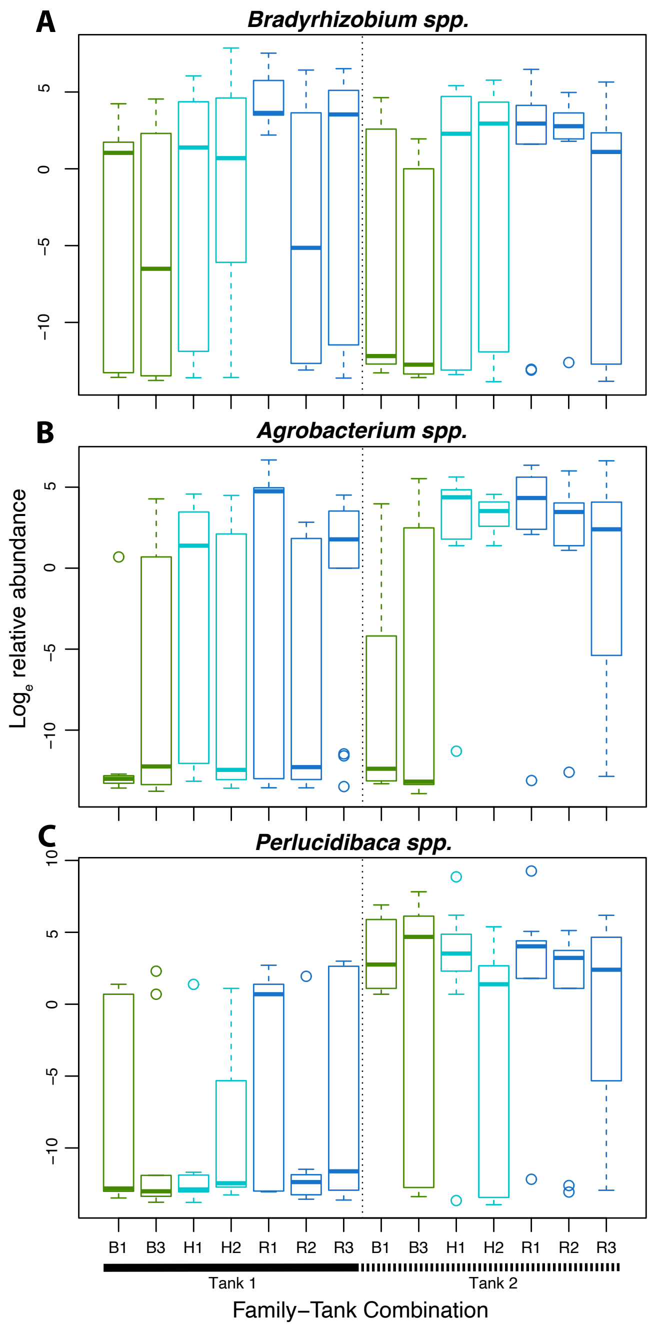


Figure S6. Relative abundances of individual ASVs differ among stickleback populations, between rearing tanks, and as a consequence of population-by-tank interaction effects. Shown are boxplots of the distributions for loge relative abundance of three ASVs: d4a5499b22ebcc23e725464ee016c371 (A), 90a96510ea0b28822ee8ccf7a323d16d (B), and 9b8bfe4d30fb93f1cd4abe8a3a7b4192 (C), with corresponding taxonomic classifications of Bradyrhizobium, Agrobacterium, and Perlucidibaca, respectively. A and B demonstrate examples of population- (genotype) by environment (tank) interaction, whereas C demonstrates an overall tank effect, on relative abundance.

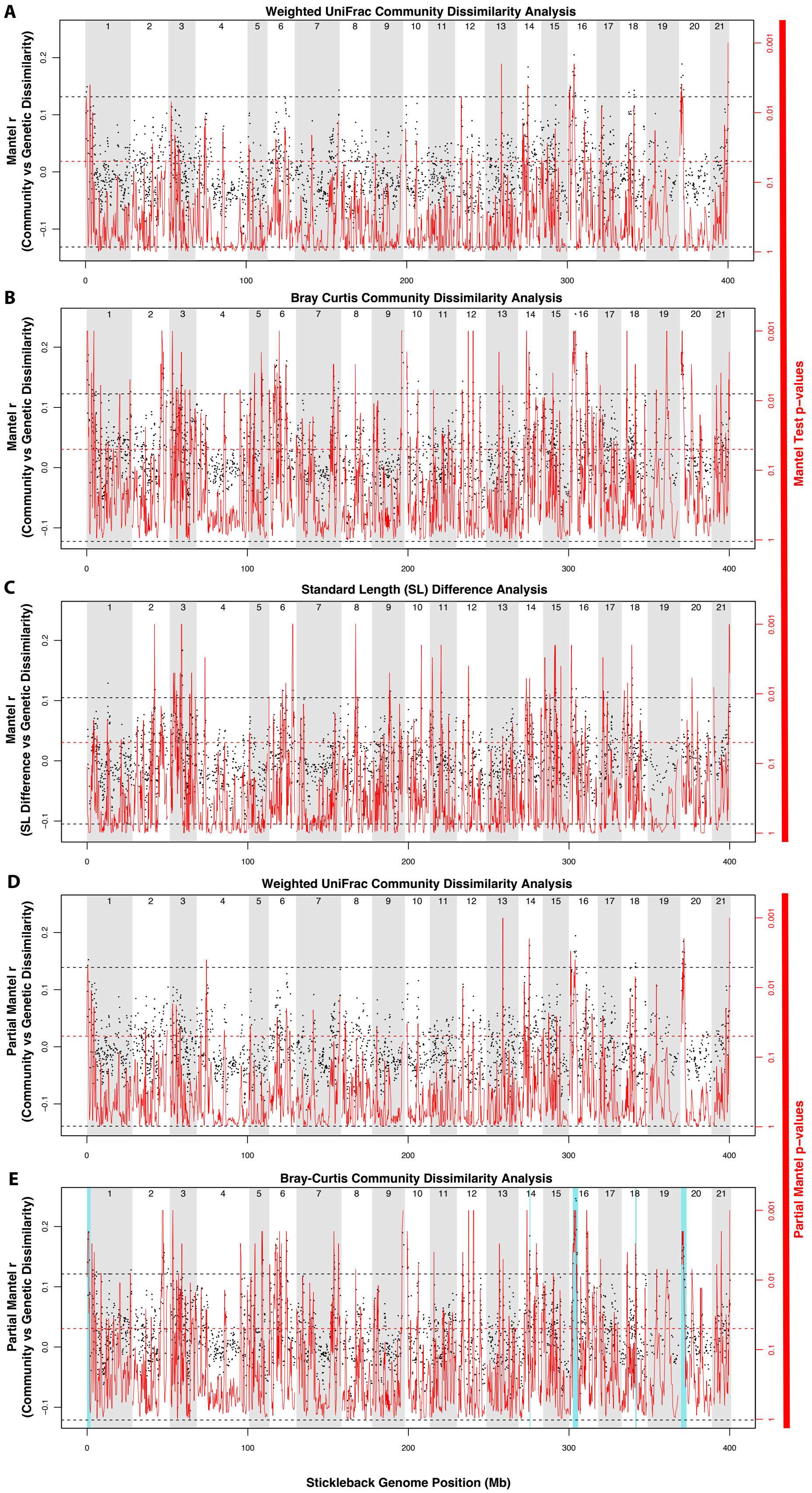

A
Weighted UniFrac Community Dissimilarity Analysis
Mantel r
(Community vs Genetic Dissimilarity)
B
Bray Curtis Community Dissimilarity Analysis
Mantel r
(Community vs Genetic Dissimilarity)
C
Standard Length (SL) Difference Analysis
Mantel r
(SL Difference vs Genetic Dissimilarity)
Mantel Test p–values
D
Weighted UniFrac Community Dissimilarity Analysis
Partial Mantel r
(Community vs Genetic Dissimilarity)
E
Bray-Curtis Community Dissimilarity Analysis
Partial Mantel r
(Community vs Genetic Dissimilarity)
Partial Mantel p–values
1 2 3 4 5 6 7 8 9 10 11 12 13 14 15 16 17 18 19 20 21
0.2 0.1 0.0 −0.1
0.001 0.01 0.1 1
0 100 200 300 400
Stickleback Genome Position (Mb)

Figure S7. Associations between host genetic dissimilarity (along the stickleback genome) and 1. Gut microbiome dissimilarity (A-B), 2. Standard Length differences (C), and 3. Gut microbiome dissimilarity after accounting for Standard Length differences (D-E). Plotted as black points along left-hand y-axes are Mantel (A-C) and partial Mantel (D-E) test statistics for the overlapping, 5-marker sliding windows. Plotted as red lines along right-hand y-axes are the respective test p-values (lowess smoothed). Note that the red p-value y-axes are on a log10 scale, ascending from high to low, to make easier comparisons with the black test statistic y-axes. In A and B, standard Mantel tests were used to measure the relationship between genetic and gut microbiome dissimilarity based on Weighted UniFrac and Bray-Curtis metrics, respectively. In C, the Mantel test reveals associations between genetic dissimilarity and size dissimilarity, as measured by the absolute value of the standard length (SL) difference between fish. In D and E partial Mantel tests were used to account for any size effects on the microbiome, although these are probably minimal based on the limited overlap of peaks between A/B and C. E is the same plot show in Fig 5A, and is included for reference here. Blue windows demarcate regions of statistical association from the Bray-Curtis analysis that overlap with those of the Weighted UniFrac analysis (D).

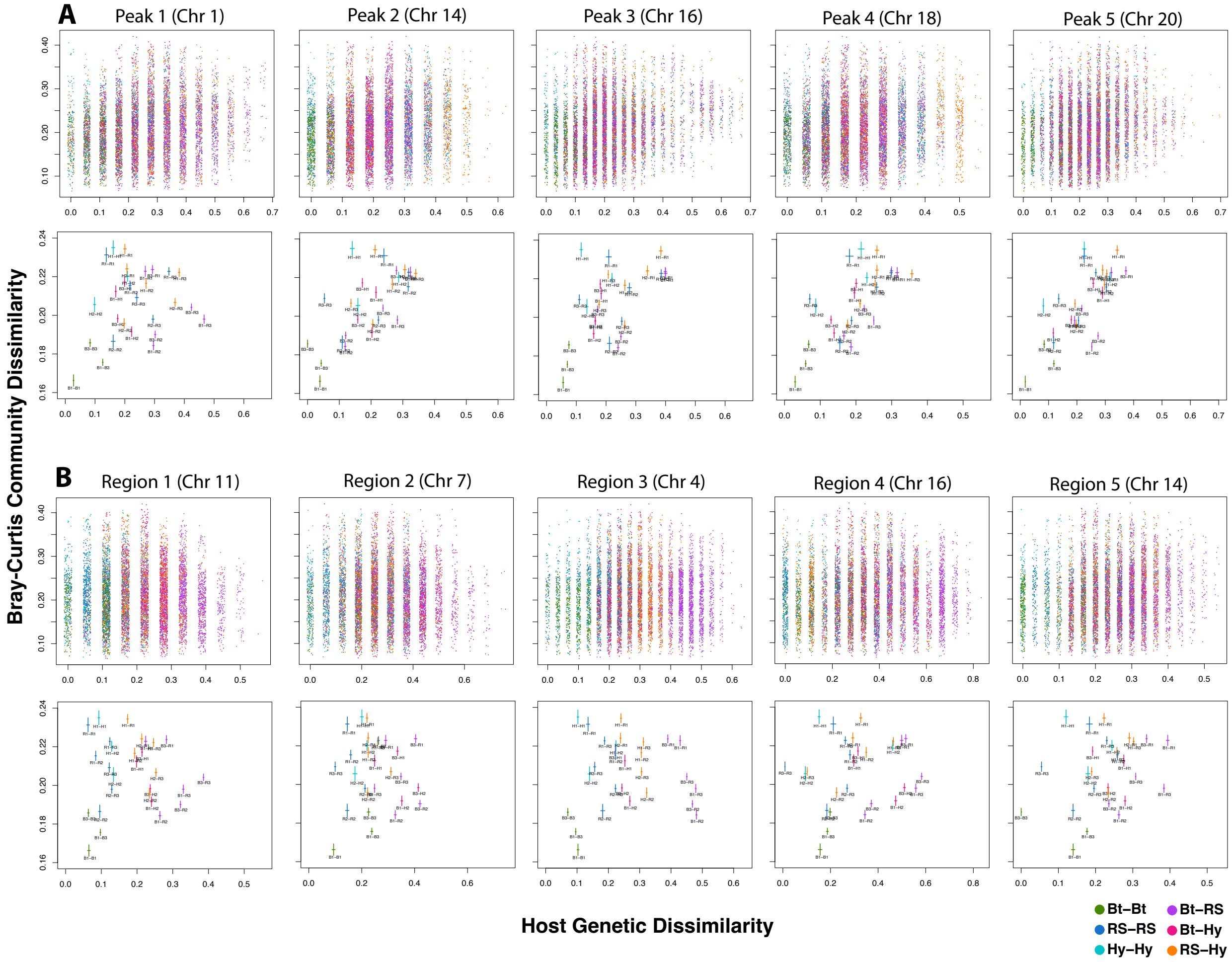


Figure S8. Differentiation at specific regions of the stickleback genome shows especially strong correspondence to gut microbiome dissimilarity, relative to randomly selected regions. Regions in A correspond to the five “peaks” of association that overlap between Bray-Curtis and

Weighted UniFrac genome-wide analyses. Regions in B were randomly selected from the genome and include the same number of markers as peaks 1-5. Plotted elements are colored (see legend) according to the population combination reflected by the fish pair. In the top row of plots for both A and B, individual pairwise dissimilarities are represented by points, which are "jittered" about discrete degrees of genetic dissimilarity (x-axis) to reduce obfuscation. In the bottom row of plots for both A and B means and means +/- standard errors are plotted as crossing lines representing family-wise stratification of the pairwise dissimilarities, with each family pair represented by a text label.

### Supplementary Sheet Captions

Sheet S1. Metadata for 149 individual stickleback fish surviving to 60 dpf during the experiment. Included is information about population and family of origin, rearing environment, standard length (SL), sex, dissector, and whether the library was excluded owing to low (< 40,000 filtered reads) sequencing coverage.

Sheet S2. Random forest (RF) classification importance metrics for 2306 ASVs. Each sub-sheet (three total) includes the mean decrease in classification accuracy observed when a given ASV was not sampled in RF iterations, relative to when it was included. The three class-specific and overall average decrease-in-accuracy values are included, along with the ASV ID and taxonomic assignment. The three sub-sheets correspond to 1. The model trained on tank 1 and tested on tank 2, 2. The model trained on tank 2 and tested on tank 1, and 3. The model trained on 80 randomly selected fish and tested on the remaining 54. ASVs highlighted in orange were consistently in the top 10 across the three models, and those in yellow were consistently in the top 10 in two of the three models.

Sheet S3. Zero-inflated negative binomial (zinb) and negative binomial (nb) tests for differential relative ASV abundance. Included are hypothesis test results for 315 ASVs with adequate sequencing coverage across samples, based on zinb models for ASVs with especially sparse distributions and nb models for ASVs with non-sparse ones (see Methods). For each term in the model tested (population, tank, standard length, and all interactions), the test statistic, p-value, and false discovery rate (FDR)-adjusted are included. The first eight columns contain the ASV IDs and assigned taxonomies.

Sheet S4. Results from a sliding window analysis of association between pairwise host genetic, microbiome, and host size dissimilarities. Each row in the sheet corresponds to an overlapping, sliding window of five adjacent genetic markers. Provided for each window are the stickleback chromosome, the locus (marker) IDs, the chromosome and whole genome coordinates for the five markers, and a series of Mantel test results. Columns 6-7 and 8-9 represent partial (accounting for SL) and standard Mantel $r$ and $p$ values for tests based on Bray-Curtis microbiome dissimilarity, columns 10-11 and 12-13 represent partial (accounting for SL) and standard Mantel $r$ and $p$ values for tests based on Weighted UniFrac microbiome dissimilarity, columns 14-15 represent standard Mantel $r$ and $p$ values for tests of association between fish genetic and body size (SL) dissimilarity. Partial Mantel test results highlighted in green mark those "significant" windows for which the $r$ value was greater than the absolute value of the genome-wide minimum *r* and the *p*-value was less than 0.05, reflecting a “beyond-noise” threshold. Marker windows highlighted in blue represent stretches of windows (based on the Bray-Curtis partial Mantel tests, the Weighted UniFrac partial Mantel tests, or both) across which there is significance overlap by proxy via shared markers. Those window stretches that are shaded in blue and include significant (green-highlighted) windows from both Bray-Curtis and Weighted UniFrac partial Mantel tests comprise the “five peaks” marked in Figures 5 and S7.

### Supplementary Methods

#### Common Garden Experimental Design

Embryos from each tank replicate (n =160) were raised in 100 x 15mm perti dishes in vacuum-sterilized embryo media until nine days post fertilization (dpf). Nonviable embryos were removed from the dishes daily and sterile media was changed every three days. Dechorionated larval fish were then moved to 9.5L tanks under “summer” conditions of 16 hours of daylight and 8 hours of night where they were fed 2mL of hatched brine shrimp naupli and fry food (Ziegler AP100 larval food) designed to mimic the diet of wild stickleback (Ziegler Bros. Inc, Gardners, PA, USA). Water temperature was maintained at 20° C with salinity at 4 ppt on a recirculating system to ensure sharing of microbial communities. At 30 dpf, fish were transferred to 20 gallon tanks under “winter” conditions (8 hours of daylight and 16 hours of night) again on a recirculating water system at 20° C with salinity at 4 ppt with diet remaining constant. This low salinity environment was chosen as previous work demonstrates that both Boot Lake and Rabbit Slough populations have high reproductive success and low mortality under these conditions while pathogen loads remain low. At 59dpf fish were starved to reduce influence of food related microbes on the microbiome profiles.

#### Dissections and Sample Preparation

At day 60 fish were prepared for dissection. To reduce stress by overstimulation, fish were placed in covered 5 gallon buckets for 30 minutes prior to dissection. Groups of 4-7 fish were then chosen randomly and placed in MS222 for euthanasia. These small groups were used to increase the accuracy of microbial profiles and reduce waiting time post-mortem until sample processing. Additionally, these groups alternated between tank replicates to allow randomization of sample processing. During the gut dissection process, fish were transferred to fresh parafilm with guts dissected alternately between two researchers with researcher 1 dissecting all odd numbered fish and researcher 2 dissecting all even numbered fish to account for researcher introduced microbes. Between each dissection, parafilm was changed and tools were washed in 95% ethanol followed by a 10% bleach solution. Fish were also sprayed in 70% ethanol prior to dissection to minimize contamination of skin microbes on gut samples.

#### DNA isolation

Gut samples were transferred to pre-weighed Next Advance sterile 1.5 mL RINO screw-cap tubes containing five-SSB32 steel beads (NextAdvance, Troy, NY, USA). Samples were then weighed to ascertain the size of each gut. Samples were then homogenized in the FastPrep (company info) for 3 rounds of 45 seconds on level 6.5 in 800uL of preheated ATL (Qiagen, Valencia, CA, USA) and 0.5uM EDTA (Boston Scientific, Marlborough, MA, USA). Aliquots of gut homogenate equivalent to 20mg of tissue were then transferred to sterile 1.5mL RINO screw-cap tubes containing 100uL of ZrOB015 zirconia beads and flash frozen in liquid nitrogen (NextAdvance, Troy, NY, USA). Samples were then homogenized again using the FastPrep (company info) for 2 rounds for 45 seconds on level 6.5 in 650uL of preheated ATL buffer (Qiagen, Valencia, CA, USA).

**Library preparation**

Amplification of the V4 16S region was performed using 1 uL of template DNA, 4.5uL of 2.79uM primer mix, and 12.5 uL NEDNext Q5 Hot Start HiFi PCR Master Mix (New England Biolabs, Ipswitch, MA, USA) using the thermo profile described by Small et al. 2019 for 22 cycles. Libraries were cleaned twice using 20 uL of Omega Mag-Bind RxnPure Plus beads and were standardized to 9.235 ng of DNA (Omega Bio-Tek Inc, Norcross, GA, USA). Pooled libraries were run in one lane of sequencing on the Illumina 4000 to obtain 150bp paired end reads.
